# Developmental DNA Methylation in the Parasitoid Wasp *Nasonia vitripennis*

**DOI:** 10.1101/2024.11.28.625666

**Authors:** CL Thomas, EB Mallon

**Affiliations:** Department of Genetics, Genomics and Cancer Sciences, University of Leicester, University Road, Leicester, UK

## Abstract

DNA methylation is a crucial epigenetic mark the development of many insect species, being essential for fertility and the progression of development in a range of organisms. However, the mechanisms underpinning the role of DNA methylation in insect development remains elusive. Furthermore, the patterns of methylation in different species can be varied. Here we aim to profile methylation across metamorphosis in the insect DNA methylation model *Nasonia vitripennis* for the first time. We find DNA methylation is at the highest in the embryo, and at the lowest in the larva. We find that the gene expression levels of *NvTet* and *NvDnmt* enzymes compliment the observed methylation patterns. Performing differential methylation analysis we find enriched GO terms for developmentally specific processes and find sites with differential methylation are share homology with developmentally linked transcription factors. Additionally, we identify sites uniquely methylated in each developmental stage, many of which also share homology with developmentally linked transcription factors. In all, we find that methylation is variable in its global methylation levels and site specific methylation throughout *Nasonia vitripennis* development, but find no obvious link with gene expression.

## Introduction

DNA methylation is an epigenetic modification where the addition of a methyl group to a cytosine can change the way in which a gene is regulated. In mammalian systems, DNA methylation is dynamic throughout with two waves of epigenetic reprogramming; the first in primordial germ cells [1] where somatic DNA methylation patterns are erased passively due to the high proliferation rate of cells, and the second after fertilisation [2] with the removal of methylation inherited from the paternal gamete with the exception of those in imprinting control regions. The maintenance methyltransferase *DNMT1*, de novo methyltransferases *DNMT3* and demethylating enzymes *TET* play crucial roles in the demethylation and remethylation during these waves.

Over the last decade, the study of insect DNA methylation has grown. In a wide range of insects, DNA methylation has proven to be a crucial driver of development, and in numerous species *Dnmt* knockouts lead to developmental or egg producing arrest [3–7]. In the silk moth *Bombyx mori*, DNA methylation has been proposed to recruit MBD2/3 to the 5’ UTR region of genes to aid with Tip60 recruitment, which in turn promotes histone H3K27 acetylation and consequentially alters gene expression [8], a mechanism proposed to influence early embryo development. Even in *Drosophila melanogaster* where there is considered to be no functional methylation, methylation levels are higher in the embryo [9], and promoter methylation of the ecdysone mediating enzyme *DmSpok* may help modulate development [10].

*Nasonia vitripennis* is emerging as one of the main insect models for studying DNA methylation. Studies to date have examined the role of DNA methylation in embryogenesis [3, 6], genomic imprinting [11], sex allocation [12], sex-biased gene expression [13], photoperiodic timing [14] and ageing [15]. Its benefits as a model include its short lifecycle, its high offspring number, its possession of both *Dnmt1* (three copies) and *Dnmt3* enzymes [16], and its relatively small genome which reduces whole genome sequencing costs.

As with other insect species, RNAi of *Dnmt1a* in *Nasonia* embryos results in developmental arrest at gastrulation [3, 6]. *NvDnmt1a* knockouts in the study of Arsala *et al*. [6] found strong reduction in gene body methylation and a corresponding reduction in gene expression. They noted that the number of methylated genes in the embryo was similar to a previous *Nasonia* study in adults [17], suggesting that whilst DNA methylation is important for embryonic viability, it is stable through development. Added to this, a different study found that reciprocal crosses maintained their parental methylation patterns through development into the F1 adult, suggesting methylation patterns remain stable through development [11]. These two findings, combined with the knowledge that RNAi of *NvDnmt1a* produces lethal effects in the F1 embryo [3, 6] but no observable phenotype through development and adulthood for the injected mother, suggest that methylation is both stable and unimportant in developmental stages following embryogenesis.

From other Hymenoptera studies, we know that global methylation levels do fluctuate [18–20], even though different species show different patterns. Consequentially, here we aim to better profile the developmental methylome of the insect DNA methylation species *Nasonia vitripennis*. We perform differential methylation analysis, and use protein binding prediction tools to help predict the role DNA methylation may have on developmental processes. By performing RNA sequencing, we are able to model the effects of DNA methylation on gene expression to attempt to identify mechanisms in which DNA methylation may regulate gene expression. Finally, we examine the expression profile of genes associated with DNA methylation to make predictions of their role in driving global methylation changes. Combined, we hope these results provide further insight to the function of DNA methylation through insect metamorphosis.

## Materials and Methods

### Sample Collection

Wasps were of the *Nasonia vitripennis* species from the Leicester strain [15]. Stocks were fed 20% sucrose solution and were maintained at 25°C with 40% humidity and 12 hour light:12 hours dark cycle.

This study focused on female *Nasonia*. Parental wasps were taken from stocks within 24 hours of eclosion, and placed three females with one male. Once removed from stocks, wasps were given 24 hours with a “practice” host placed in a cut pipette tip pierced through a cotton bung (for ease of egg collection), and filter paper soaked with 20% sucrose. The practice host was then replaced with a fresh host, and the wasps were given three hours to lay. After three hours, the host was removed from the parental wasps and incubated until the appropriate collection time. This produced 83.3% females (n = 944), similar to previous developmental studies that used a single female (80-95%) [17, 21]. The following samples were collected; Embryo (7-10 hours post laying), larvae (69-72 hours post laying), prepupa (7 days post laying), pupa (yellow eye pupa 9 days post laying), and adult (24 hours post eclosion). Adults and pupa were sexed by inspection.

DNA for whole genome bisulphite sequencing was extracted using the DNeasy Blood & Tissue Kit (Quiagen) with some adjustments to the standard protocol (see [22]).

Larval samples were found to be RNA rich, so RNase incubation was increased to 90 minutes. 1,700 embryos, 30 larvae, 10 prepupa, 10 pupa and 10 adults were used for each replicate of the developmental stage samples. Three replicates were used for each stage with the exception of the embryo where one sample failed quality control. Quality of DNA samples was tested using a NanoDrop 2000 spectrophotometer (Thermo Scientific), a QubitTM dsDNA BR Assay Kit (ThermoFisher Scientific) and by running a 1% agarose gel for 40 minutes at 100V. DNA samples were sent to BGI Genomics for bisulphite treatment, lambda spiking, adapter ligation, sequencing and trimming.

For RNA samples 400 embryos, 30 larvae, 10 prepupa, 10 pupa and 10 adults were used per replicate (3 replicates for each stage). Samples were extracted using the Tri-Reagent (Sigma Aldrich) protocol, with adjustments for TURBO DNase ^TM^ treatment and removal (as outlined in [22]). RNA concentration was measured using the QubitTM RNA BR Assay Kit (ThermoFisher) and the integrity of RNA was assessed on a Agilent 2100 Bioanalyzer using the RNA 6000 Nano Kit (Agilent). RNA samples were sent to BGI Genomics for sequencing and trimming.

### Bioinformatics

#### DNA Methylation

To produce an overview of the number of methylated sites at each developmental stage, all replicates for each stage were merged to create one fastq file for each developmental stage. Each fastq file was aligned to the N Vit 1.1 PSR genome [23] using Bismark [24]. Samples were also aligned to the lambda genome to test the percentage of false positives. The Bismark deduplicate command was used to remove PCR replicates, and the inbuilt Bismark bismark methylation extractor command [24] was used to extract the methylation level. Methylation status was destranded to improve coverage using the inbuilt Bismark coverage2cytosine command [24]. A binomial test was performed for each developmental stage to determine the methylation status of each site on R (version 4.3.1), where probability was set as the mean lambda alignment percentage (FDR *<* 0.05).

For the differential analyses, each replicate was aligned, deduplicated, extracted, destranded and tested for methylation status as outlined above. Samples were filtered using the R package methylkit [25] so that only CpGs with more than 10 read coverage were analyses. A PCA was generated using the MethylKit PCASamples command, and four pairwise comparisons were established using the MethylKit reorganise command. These comparisons were embryo versus larva, larva versus prepupa, prepupa versus pupa and pupa versus adult. Differentially methylated CpGs were those with a statistically significant difference (P *<* 0.0125 (Bonferroni Correction)) and a methylation difference greater than 10%.

Genomic features were then assigned to the methylated sites using a GFF file containing all the genomic features created using AGAT [26]. Transcription factor binding sites were established using TFBSTools [27], using the JASPAR 2024 [28] *Drosophila melanogaster* transcription factor list, identifying matches with 100% homology. To identify enriched genomic motifs HOMER was used [29] with the -size paramater set as 20. GO terms were obtained from the Hymenoptera Genome Database [30], and enrichment was performed using the R package TopGO [31].

Whole genome bisulphite sequencing files were made available on EBI with the accession number: E-MTAB-14656.

#### RNA

Paired-end RNA reads were aligned to the Nvit PSR1.1 genome [23] using the Star (Version 2.7.5c) 2pass method [32]. Read count number was generated using HTSeq [33] and normalised using DESeq2 (1.32.0) [34]. For comparison of expression levels for genes associated with methylation, the DESeq2 fragments per million mapped were generated using the fpm() command and plotted using ggplot2 [35]. Exon counts were generated by DEXSeq [36]. RNA libraries are available on EBI under the accession number: E-MTAB-14657.

#### Modelling Methylation with Gene Expression

Generalised linear models (GLMs) were used to analyse the effects of DNA methylation levels on gene expression. Whole gene methylation effects on gene expression were analysed with a GLM with a quasipoisson distribution with the model: RNA Counts ∼ Methylation Percentage * Developmental Stage. When looking at methylation effects on exon usage we used a GLM with a quassipoisson distribution with the model Exon Counts ∼ Methylation Percentage * Developmental Stage. Next, we categorised genes as highly methylated (70-100% methylated), medium (30-70%), low(0.5-30%), or not methylated (*<* 0.5%) as previous insect developmental studies had found differences in gene methylation categories between different developmental stages [20]. With these categories we examined the effect these had on gene expression using a quasipoisson distribution with the model; Gene Counts ∼ Methylation Category * Developmental Stage.

We analysed the effect of the number of differentially methylated sites within the gene on gene expression using a GLM with a gaussian distribution (which deals with both positive and negative values) with the model log fold change ∼ Number of differentially methylated sites * pairwise comparison type. We analysed the effect of differentially methylated sites within predicted transcription factor binding sites on gene expression with a GLM with a gaussian distribution using the model gene log fold change ∼ distance to start site * transcription factor * pairwise comparison. Finally, when examining if the number of consecutive differentially methylated CpGs influences gene expression we used a GLM with a gaussian distribution with the model; log2lfc distance to TSS * pairwise comparison * consecutive number of CpGs.

The script for all annotations, methylation processing and RNA count is available on github at:

https://github.com/C-L-Thomas/Nasonia_Developmental_Methylation.

## Results

### Developmental Stage Methylation

Of the 16,515,404 destranded CpG sites in the *Nasonia vitripennis* genome, between 14,903,835 and 15,084,779 had a coverage of at least 10 when combining all reads for each developmental stage. We define a methylated site as CpG sites that pass a binomial test with the false positive rate calculated using a lambda spike for each bisulphite treated sample. The number of destranded methylated sites across each developmental stage was relatively stable; 182,740 for embryo (1.2% of CpGs), 170,115 for larva (1.1% of CpGs), 178,611 for prepupa (1.2% of CpGs), 182,221 for pupa (1.2% of CpGs) and 178,309 for adult (1.2% of CpGs).

The embryo had the highest percentage methylation for these methylated CpGs (**Figure 1A**), before a drop in the percentage methylation into the larval stage. The percentage methylation increased slightly into the prepupal larva stage before remaining reasonably stable for the remaining developmental stages. The location of methylated CpGs was similar in each developmental stage (**Figure 1B**), with most methylation being in the gene body, predominantly in the exon.

**Figure 1.**
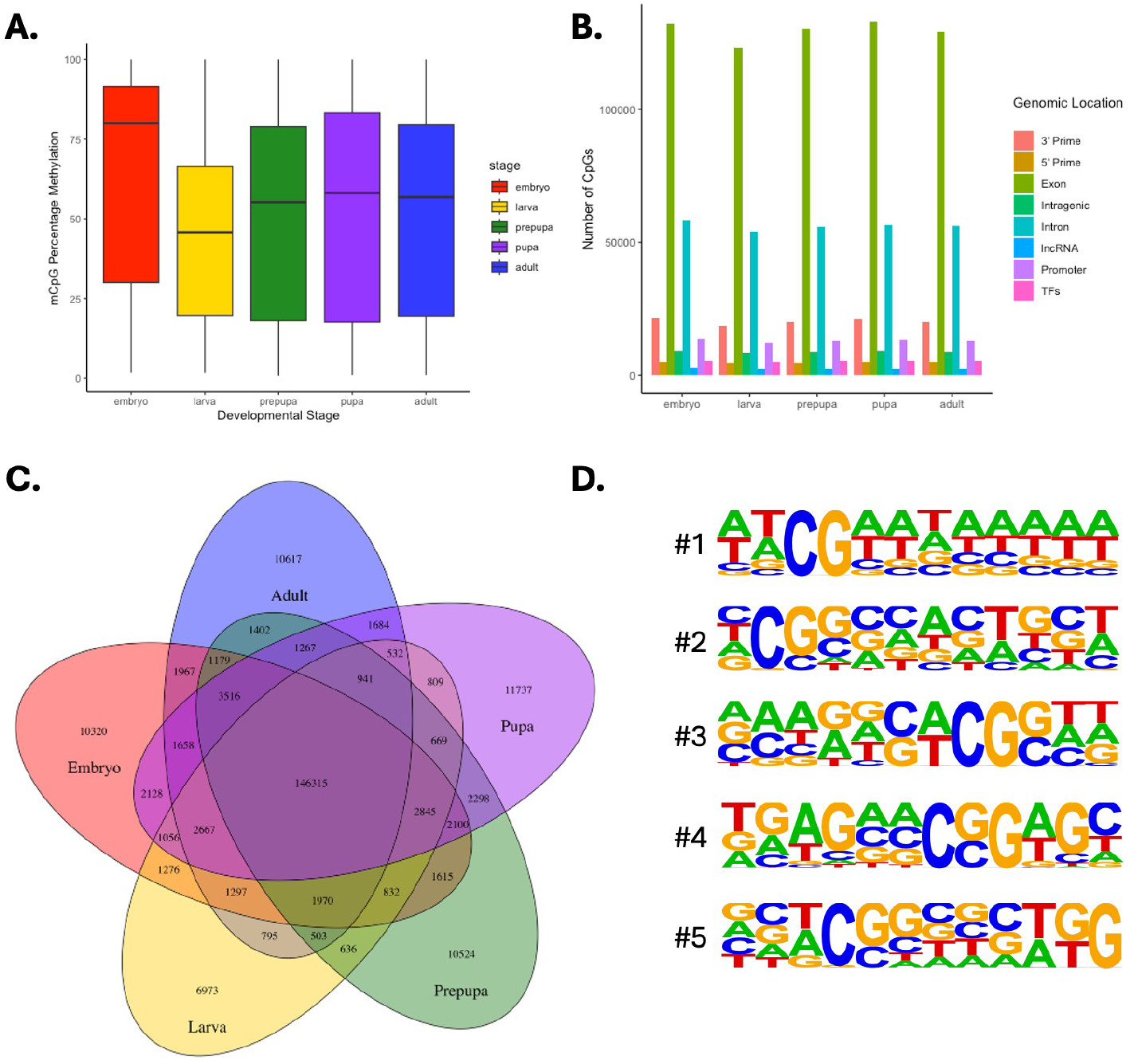
Developmental Stage Methylation. **A**. Percentage methylation for CpGs that pass binomial test (ie. are methylated). **B**. Genomic location of methylated sites for each developmental stage. **C**. Venn diagram showing the number of CpGs that are methylated between different developmental stages. **D**. Top five methylated motifs of the sites that are methylated in all five developmental stages.

146,315 CpGs were commonly methylated in all developmental stage (Figure 1C), significantly fewer than expected by chance (*χ*^2^ = 1787302, df = 1, p *<*2.2 *×* 10^−16^). The most enriched motifs of these shared CpGs throughout development are found in **Figure 1D**. The methylated CpGs found in all stages had protein binding motifs with sequence homology to that of *Drosophila* DEAF-1, runt, dref, zeste, brinker, odd-paired, mothers against dpp and giant protein binding sites. Amongst the CpGs methylated only in the embryonic stage, we found motif enrichment for sites with homology to the *Drosophila* BEAF-32, smooth and Abd-A binding sites. In the larva, we found enrichment for binding site with homology of the biding sites for the segmentation transcription factors odd and buttonhead, as well as the transcription factors ovo and suppressor of hairless. In the sites uniquely methylated in the prepupa, we found enrichment for motifs that share sequence homology with *Drosophila* brachyenteron, pointed, grainy head, and pangolin transcription factors. In sites uniquely methylated in the pupa, we found motifs that share homology with *Drosophila* sine oculis, kruppel and ventral veins lacking transcription factors. In the adult, we find transcription factor binding motifs with similarities to *Drosophila* snail, Transformer-2 sex-determining protein, and vismay.

### Differential Methylation

Next we sought to identify differentially methylated CpGs. At a CpG level, replicates from each developmental stage clustered together in a PCA (**Figure 2A**). To examine the trajectory of methylation between different developmental stages, we performed four pairwise differential analyses progressing through the four developmental stages. In these analyses, the majority of GO terms of genes that possessed differentially methylated CpGs were shared across all four comparisons. As a consequence, we only highlight GO terms that had more significant enrichment for each of the pairwise analyses. Some examples of highly enriched terms in all analyses include multicellular organism development, imaginal disc-derived wing morphogenesis, oogenesis, cell differentiation, anatomical structure morphogenesis and axonogenesis. The genomic location of the differential expression for each developmental stage (ie. exon, intron, intragenic) followed a similar pattern to that seen in **Figure 1**, indicating that differential methylation had no preference for particular genomic location.

**Figure 2.**
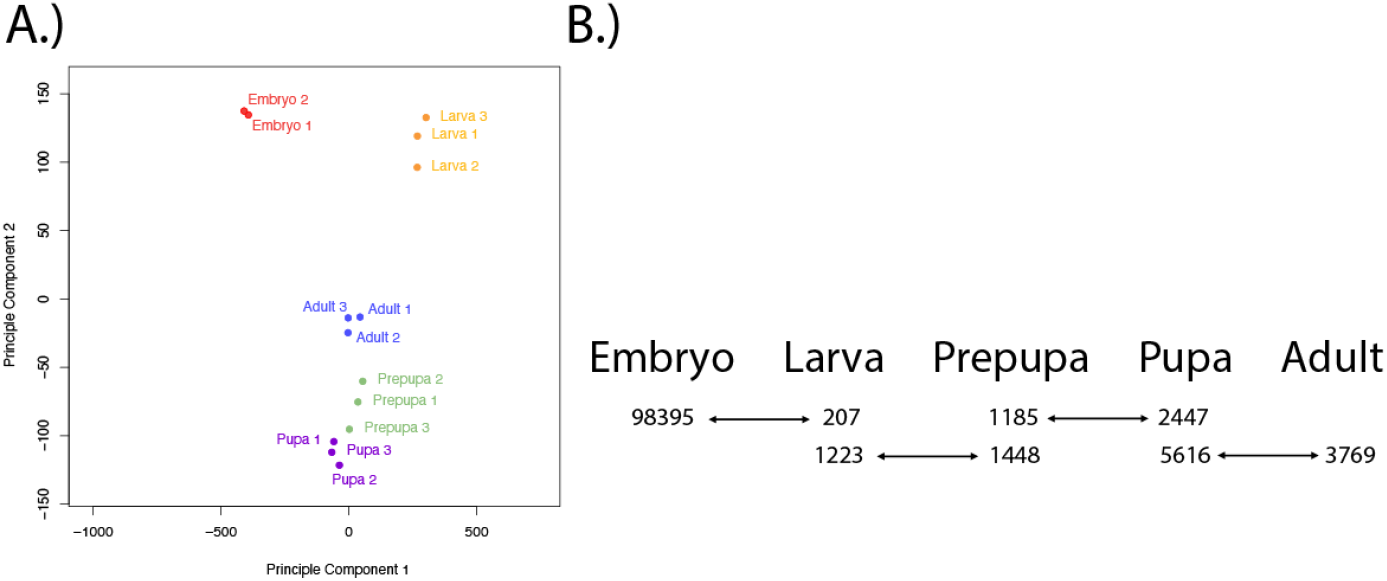
Differential Expression Results. **A**. PCA of each individual whole genome bisulphite sequencing sample. **B**. Schematic of each pairwise comparison, with the number of differentially methylated CpGs

Most differential CpG methylation occurs between the embryonic and larval stages of development, with 98,602 differentially methylated CpGs; 98,395 hypermethylated in the embryo, 207 in the larva (**Figure 2B**). The genes that these differentially methylated CpGs were located in had biological process GO terms cell development (GO:0048468), embryo development (GO:0009790), pattern specification process (GO:0007389), gliogenesis (GO:0042063) and methylation (GO:0032259). Running motif analysis on the differentially methylated sites, we found protein binding sites similar to those of extradenticle, deaf1 and brachyenteron.

Between the larval and prepupal stages of development there were 15,718 differentially methylated CpGs, the majority of which were hypermethylated in the prepupa (**Figure 2B**). GO term analysis revealed genes enriched for carbohydrate metabolic process (GO:0005975) and R3/R4 cell differentiation (GO:0048056). Running motif analysis on the differentially methylated sites, we found enrichment for *Drosophila* glass, deaf1, and pangolin binding sites.

Between the prepupa and pupa stages of development there were 3632 differentially methylated CpGs, 2447 hypermethylated in the pupa, 1185 in the prepupa. GO term analysis revealed a unique set of enriched terms for cell fate commitment (GO:0045165), regulation of developmental growth (GO:0048638), and neurotransmitter secretion (GO:0007269). Running motif analysis on the differentially methylated sites, we found motifs enriched for Mothers against dpp, Adh transcription factor 1, ap1, kruppel, serpent, vismay, bagpipe, biniou and Pleiohomeotic polycomb protein binding.

Between the pupal and adult stages of development we found 9385 differentially methylated CpGs, 5616 hypermethylated in the pupa, 3769 hypermethylated in the adult. Unique GO terms found in the pupal adult pairwise analysis included animal organ development (GO:0048513), neuron projection development (GO:0031175), morphogenesis of an epithelium (GO:0002009), and neuron fate commitment (GO:0048663). Running motif analysis on the differentially methylated sites, we found motifs enriched for tramtrack binding, buttonhead, ventral nervous system defective, chorion factor 2, giant, ovo and trithorax-like.

Whole gene differential methylation levels can be found in the **Supplementary Information 1**.

### DNA Methylation’s Effect on Gene Expression

To better understand the link between DNA methylation and gene expression, we collected RNA from each developmental stage samples. We found the demethylase enzyme *TET* has highest expression in the embryonic stage of development (**Figure 3**), and its drop in expression coincides with the biggest demethylation event observed between the embryonic and larval stages of development (**Figures 1A & 2B)**. We also observed *Dnmt1a* expression highest in the embryo, remaining the *Dnmt* enzyme with highest expression throughout development. We see expression of *Dnmt1c* and *Dnmt3* remain at similar levels until the adult stage, where *Dnmt1c* is expressed at a similar level to *Dnmt3*. This is consistent with the theory that *Dnmt1c* may be an ovary specific methyltransferase [22]. We found no expression of *Dnmt1b*.

**Figure 3.**
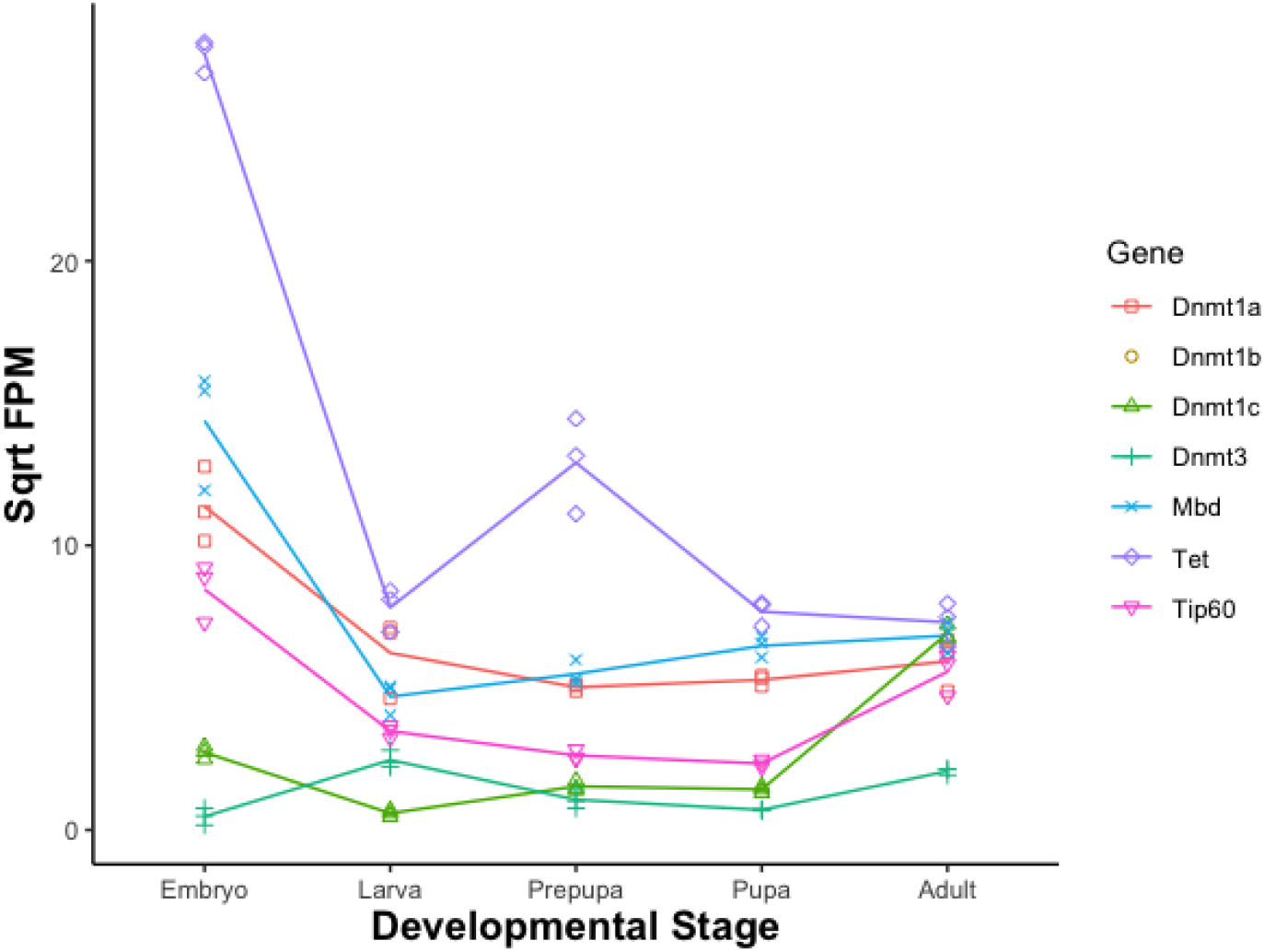
Square root fragments per million mapped fragments of genes associated with DNA methylation

To investigate whether a relationship existed between DNA methylation and the regulation of gene expression we ran multiple GLMs. The first GLMs used a quasipoisson distribution and investigated the relationship between gene level methylation and gene expression (**Table 1A** & **Figure 4A**), exon methylation and exon count (**Table 1**,& **Figure 4B**) and gene methylation category (none, low, medium, or high) and gene expression (**Table 1C** & **Figure 4C**), however all models had a poor model fit. Next we examined how the differential methylation effects changes in gene expression between developmental stages. Using a gaussian distributed GLMs we investigated the effect of number of differentially methylated sites within a gene on gene expression (**Table 2A**) and the effect of differentially methylated CpGs within transcription factor binding sites, and their distance to their closest gene transcriptional start site (**Table 2B** & **Figure 4D**), but both of these had a low pseudo-R^2^.

**Table 1.** Quasipoisson Distributed GLMs.

| Analysis | Model | Chisq | df | p-val | Pseudo R<sup>2</sup> |
| --- | --- | --- | --- | --- | --- |
| A. | Gene RNA Counts ~Gene Meth % * Dev Stage | 40.107 | 4 | 4.1e-8 | 2.08 |
| B. | Exon RNA Counts ~Exon Meth % * Dev Stage | 9.9 | 4 | 0.04275 | 1.2 |
| C. | Gene RNA Counts ~ Meth Category * Dev Stage | 39.8 | 12 | 7.8e-5 | 2.3 |

**Table 2.** Gaussian Distributed GLMs.

| Analysis | Model | Chisq | df | p-val | Pseudo R<sup>2</sup> |
| --- | --- | --- | --- | --- | --- |
| A. | log2lfc~DMS*stage | 100.72 | 3 | 2.2-e16 | 2.1 |
| B. | log2lfc~disttss*stage*TF | 11.27 | 27 | 1 | 5.3 |
| C. | log2lfc~distance*stage*consecutive | 0.083 | 2 | 0.9593 | 26 |

**Figure 4.**
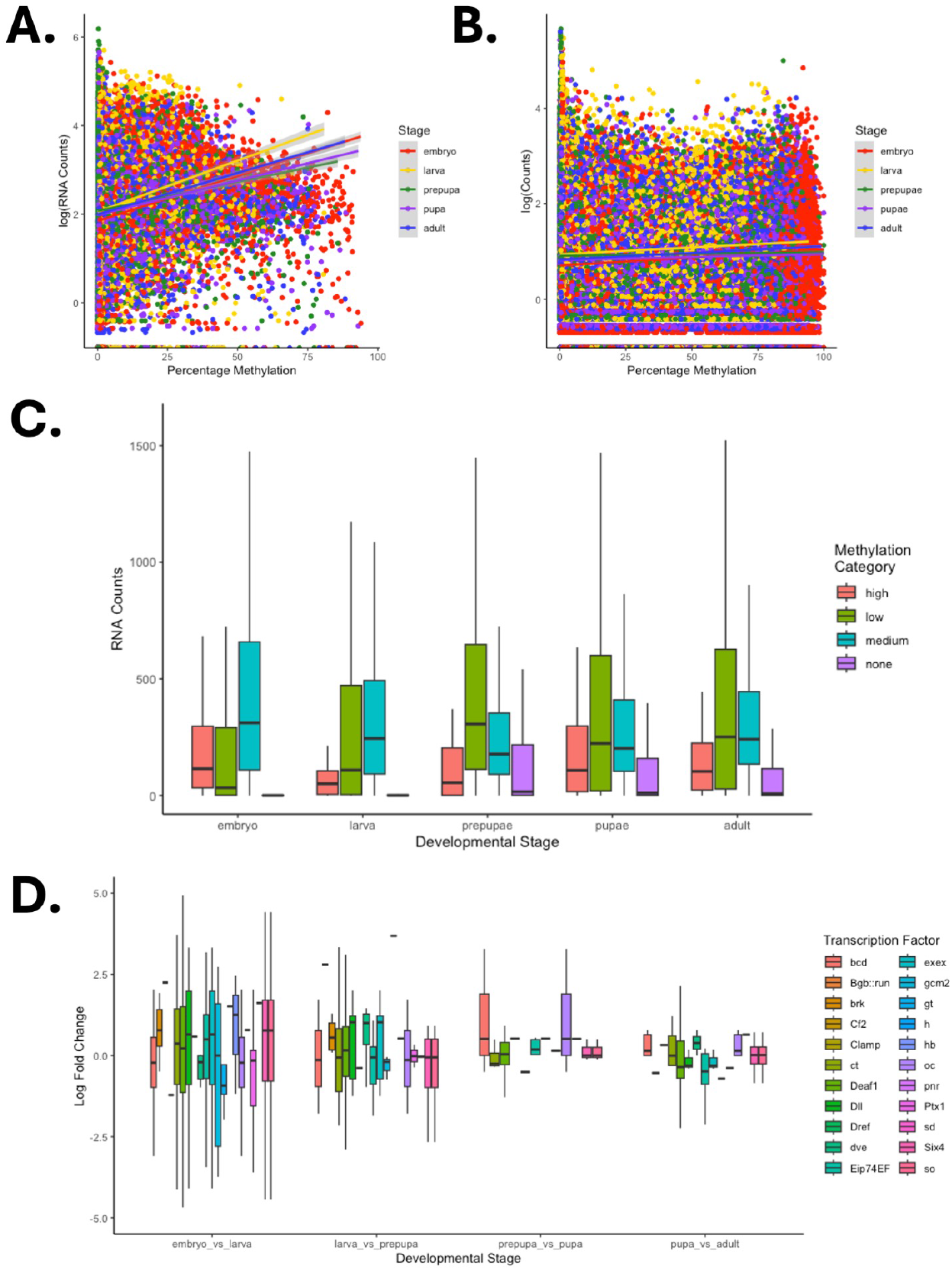
Linking DNA methylation with gene expression. **A**. Gene level methylation against gene RNA counts. **B**. Exon level methylation against exon RNA counts. **C**. Gene methylation category against average RNA gene count. **D**. The log fold change in gene expression for transcription factors between each developmental stage

Finally, we investigated whether the number of consecutive differentially methylated CpGs influenced gene expression levels of the closest gene, as the study by Xu *et al*. [8] indicated that more methylated CpGs had a greater effect on gene expression. By selecting genomic locations that have two or more differentially methylated sites consecutively we were able to generate a model with better fit (26%, **Table 2C**), but still not a good enough fit to make interpretations. However, a point of interest is that while the model had a pseudo-R^2^ of 25.5% if run only for sites with two consecutive differentially methylated CpGs, this improves to a pseudo-R^2^ of 46% for sites with three consecutive differentially methylated CpGs. Full tables of these GLMs can be found in **Supplementary Information 3**.

## Discussion

In this study, we examined DNA methylation levels through different stages of metamorphosis of the DNA methylation insect model *Nasonia vitripennis*. We find variable levels of methylation between developmental stages (**Figure 1A**)). Despite this, the majority of methylated CpGs remain methylated across all developmental stages (**Figure 1C**)). We ran motif analysis on the sites which were commonly methylated in all developmental stages and found enrichment for sites sharing sequence homology with important developmental transcription factor binding sites in *Drosophila melanogaster* including the multi-developmental process regulating runt [37] and dref [38] proteins, the decapentaplegic regulators brinker [39] and mothers against dpp, and other developmental proteins including giant, odd-pair and Deaf1. Stage dependent, we find methylation in 1.1-1.2 % of CpGs, which places our findings in the middle of the 0.63% found by Beeler *et al*. [40] and the 1.6% found by Wang *et al*. [17]. Similar to the study of Wang *et al*. [17] we find methylation predominantly in the gene body (**Figure 1B**)), particularly in exons towards the 5’ end. As exon number increases, methylation level decreases.

We observed the highest methylation levels in the embryonic stage (**Figure 1A**)). Following the embryonic stage, we observe methylation at its lowest level in the larval stage (**Figure 1A**)). The difference between these two stages is evidenced in the differential site methylation analyses where most differential methylation occurs between the embryonic and larval stages (**Figure 2B**)). This pattern fits with the results of a dot blot assay in a previous silk moth study [7] where methylation levels drop throughout embryogenesis following gastrulation. Methylation levels increase as the larva prepares for the pupal stage of metamorphosis (**Figure 1A**). Following the prepupal stage the global level of methylation remains relatively stable (**Figure 1A**), with increased local fluctuations in site methylation (**Figure 2B**). The patterns observed in this study are similar to those observed in the honey bee [18] where methylation is also highest in the embryo, lowest in the embryo, with later developmental stages in between these two states. However, both the honeybee and *Nasonia* results here contrast another Hymenoptera species the Florida carpenter ant *Camponotus floridanus* where the embryonic stage has the fewest methylated sties [19].

Between the developmental stages we see hallmarks suggesting that DNA plays a role in many key developmental processes. For example, between the embryo and larval stages of development we find GO terms embryo development, pattern specification process, gliogenesis, and methylation. As the wasp enters the larval stage of development we find CpGs uniquely methylated at this stage that share homology with the binding sites for *Drosophila* buttonhead and the odd-skipped transcription factors, which have both previously been implicated in larval neurodevelopment [41, 42]. In the transition between the larval and prepupal stages we find differentially methylated CpGs in genes with GO term enrichment for R3/R4 cell differentiation, which is fitting for this developmental stage as this is an important step in establishing planar cell polarity in the *Drosophila* eye [43]. We also find enrichment of differentially methylated CpGs for sites that share homology with the Drosophila external sensory organ protein Musashi [44] and the larval development protein pangolin. Once the developing wasp enters the prepupal stage, we find that sites only methylated in this stage share sequence homology with the*Drosophila* wing development proteins pointed and grainy head [45]. Transitioning from the prepupal stage into the pupa we find differentially methylated CpGs located in genes enriched for developmental growth, cell fate commitment and neurotransmitter secretion, as well as motifs which share homology with the *Drosophila* vestigial wing-patterning protein Mothers against dpp [46], the adult sense organ precursor cell regulator Suppressor of Hairless [47], and the well known patterning transcription factor kruppel. In the final transition as the wasp goes from the pupal stage and ecloses as an adult, we find differentially methylated CpGs located in genes enriched for GO terms including animal organ development, neuron projection development, morphogenesis of an epithelium and neuron fate commitment. We also find motif enrichment of these differentially expressed CpGs that share homology with the *Drosophila* neurogenesis protein ventral nervous system defective binding site. With our results giving stage specific GO terms and finding enrichment of motif binding for protein binding sites with stage associated proteins, it appears that despite *NvDnmt1a* knockouts only being lethal in the embryo [3, 6] methylation continues to play a role in regulating stage specific processes.

We also performed RNA sequencing to examine the link between DNA methylation and gene expression. Given that *DNMT* s and *TET* are crucial drivers of mammalian [48] and insect [3, 6] development, we decided to examine how their expression through development compares with the developmental fluctuations in DNA methylation (**Figure 3**). We see *NvTet* has the highest expression in the embryo, which pairs with the biggest drop in methylation between the embryonic and larval stages of development. This posits the question of the importance of *NvTet* in development. As far as we’re aware, there is no literature for *Tet* RNAi for in insects. It would be interesting to see the importance of this enzyme, and see if it’s knockout produces a similar arrest at gastrulation as *NvDnmt1a* does [6], or whether a later developmental arrest between the embryonic and larval stages occurs. Perhaps *NvTet* is not essential for development, and methylation between the embryo and larval stage is lost passively by high cell replication rate. However, the results here suggest a role of *NvTet* in the transition from the embryo to the larva. Given the vital role of DNA methylation in *Nasonia* embryogenesis, it is also worth noting that *NvMbd* and *NvTip60*, the only proven insect methylation effectors [8], have their highest expression in the embryonic stage of development. A final observation regarding methylation associated genes is that unlike each other methylation associated protein, *NvDnmt3* has increased expression between the embryo and the larval stage. Given the increase in methylation between the larval and prepupal stages (**Figure 1A** & **Figure 2B**), this hints at a de novo methyltransferase role for *NvDnmt3* similar to that in mammals, as increased expression may allow targeting of methylation to new genomic sites, for example sites such as pangolin, brachyenteron, and grainy head.

Although we observe a positive relationship between gene (and exon) methylation with gene (or exon) expression (**Figure 4A** & **B**), this is caveated with the poor model fit. Interestingly, we see genes with medium levels of methylation have a higher average gene expression level in the embryo and larva, whilst lowly methylated genes have higher expression in the latter stages (**Figure 4C**). This may hint at a mechanism where higher gene methylation is crucial for embryo viability (as proven by Arsala *et al*. [6], but this mechanism of methylation assisting methylation in early development ends following the demethylation event in the larval stage. If true, this could in part be due to the depleting levels of the developmentally implicated *NvTip60* and *NvMbd* enzymes [8] (**Figure 3**). Finally, when focusing on consecutive differentially methylated genes, we observed better model fit when correlating gene expression with methylation loci with three consecutive differentially methylated CpGs compared to two. Whilst both models had a poor fit, this lends evidence to the finding that an increased number of consecutive CpGs has an effect on gene expression [8].

## Conclusions

Here we show the developmental methylome of the insect methylation model *Nasonia vitripennis* for the first time. We find global fluctuations in DNA methylation levels, with methylation at its highest level in the embryo, and its lowest level in the larva. Global methylation levels remain relatively stable for later developmental stages, but we still see loci differences. We performed differential methylation analysis and found GO terms and transcription factor binding motifs enriched for processes specific for each stage’s transition. We find links between the expression profiles of methylation associated genes and the global methylation, but fail to identify any further discernible links between DNA methylation and gene expression.

## Supporting Information

**Supplementary Information 1:** Results from DNA methylation analyses

**Supplementary Information 2:** Results from RNA analyses

**Supplementary Information 3:** Statistical results from GLMs linking DNA methylation to gene expression.

## Acknowledgments

CT was supported by a BBSRC MIBTP DTP studentships. EM was funded by BBSRC Pioneer Awards APP3335 – Does sleep disruption affect epigenetic ageing in an insect model?

